# AAV-delivered CRISPR-Cas9 elicits persistent retinal immune responses compared with transient responses to RNP

**DOI:** 10.64898/2025.12.11.693665

**Authors:** Juliette Pulman, Duohao Ren, Laura Visticot, Hugo Malki, Yeqian Yao, Anne De Cian, Divya Ail, Jean-Paul Concordet, Deniz Dalkara, Sylvain Fisson

## Abstract

CRISPR-Cas9 is a powerful gene-editing tool with great potential for treating genetic diseases, including inherited retinal disorders. However, its bacterial origin can induce immune responses that may eliminate transduced cells, threatening editing efficiency. A deeper understanding of CRISPR-Cas9 immunogenicity is therefore needed. Previous studies have shown that systemic delivery via Cas9 induces an immune response, but the detailed inflammation and the impact of the vector remain unclear, especially in immune-privileged organs like the eye. In this study, we found that Cas9 delivered to the retina using adeno-associated virus (AAV) induced persistent inflammation, whereas delivery as naked ribonucleoprotein (RNP) complexes resulted in acute inflammation that faded three weeks post-injection. Inflammation was more severe in the rd10 mouse model of inherited retinal degeneration, which exhibits basal inflammation. These findings provide new insights into vector-dependent immune responses to Cas9 in the eye and highlight potential risks associated with its clinical application.

**TEASER:** Understanding immune reactions to CRISPR-Cas9 and linking these to their delivery methodology increases their safety.

## INTRODUCTION

The CRISPR-Cas9 system, composed of Cas9 protein and single-guide RNA (sgRNA), enables precise, efficient, and versatile gene editing (*1*) and offers new possibilities for treating various genetic diseases (*2–4*). Despite these promises, Cas9 carries the risk of being recognized as a foreign antigen by the mammalian immune system and presented by antigen-presenting cells to lymphocytes, inducing an immune response due to its bacterial origin (*5*). This represents a risk for *in vivo* applications of CRISPR-Cas9 in gene editing. Indeed, it has been reported that Cas9 induces immune responses in preclinical gene therapy research. A previous study showed that Cas9 delivery to the liver induces CD4⁺ and CD8⁺ T-cell infiltration and anti-Cas9 antibody production in a murine model, with transduced cells being cleared within a few weeks after injection(*6*). Another study in canine muscular dystrophy models also showed that a Cas9-specific immune response was induced, including local CD4^+^ and CD8^+^ T-cell infiltration in injected muscle and the production of anti-SpCas9 antibodies. Cas9 can also induce activation of human monocytes through the STING-STAT6 pathway (*7*), and Cas9-derived peptides are presented by MHC II to elicit proliferation of CD4^+^ T-cells (*8*). Finally, in a canine model, the Cas9-edited cells were eliminated weeks after injection, even under immunosuppression with a high dose of prednisolone (*9*).

The eye is considered an immune privileged organ but this does not entirely eliminate inflammatory risk. Indeed, viral vectors, such as adeno-associated virus (AAV), can induce local inflammation and T-cell infiltration into the retina after intraocular delivery (*10–13*), adding to concerns about Cas9-induced inflammation (*5*, *9*, *10*, *14*). In the eye, EDIT-101, an AAV encoded CRISPR-Cas9 based gene therapy, was developed to treat CEP290-associated inherited retinal degeneration. A phase I clinical trial evaluated the safety profile of the therapy in 14 patients (NCT03872479). Ocular adverse events occurred in 50% of participants, leading to visual impairment in two patients, even though they were under prednisone immunosuppression. It remains unclear whether the immune response was directed against the AAV vector or Cas9 itself (*15*). To our knowledge, the immune response specific to CRISPR-Cas9 in the retina is still unclear and warrants further exploration.

Moreover, the use of viral vectors to deliver Cas9 results in long-term expression of the Cas9 transgene thereby leading to genotoxicity by increasing off-target activity (*3*). Transient delivery of CRISPR-Cas9 can be an alternative to avoid these issues by directly delivering the Cas9 protein and its guide RNA complexed as a ribonucleoprotein (RNP) (*3*). A CRISPR-Cas9 RNP complex allowing immediate genome-editing activity upon delivery and is degraded after a few days. However, this approach can also result in Cas9 presence in the extracellular matrix, increasing the risk of immune cell recognition (*16*). Generally speaking, the impacts of different delivery methods on immune responses triggered by Cas9 gene editing are currently understudied and hold important insights into the efficacy and safety of gene editing in the retina.

Furthermore, retinas from patients with inherited retinal dystrophies (IRDs) often exhibit photoreceptor degeneration, potentially leading to local microenvironmental inflammation (*17*), a phenomenon also observed in animal models of retinal degeneration (*18*). This pre-existing inflammation may exacerbate immune responses against the gene therapy, posing another risk to its safety and efficacy (*19*).

In this study, we aimed to understand the immune response induced by CRISPR-Cas9 in the retina of wild-type mice and to compare the immune consequences of different subretinal delivery methods, including direct RNP delivery or delivery of CRISPR-Cas9 DNA packaged into an AAV. Additionally, we monitored immune consequences in a retinal degeneration model (rd10 mice), representing the pre-existing inflammatory microenvironment in patients. Our results reveal that Cas9 encoded via AAV elicits a persistent immune response, activating both innate and adaptive immune markers, whereas Cas9-RNP results in a transient immune response. As expected, the immune response induced by CRISPR-Cas9 is more severe in the pathological rd10 mouse model. We demonstrate that the delivery method affects the safety and efficacy of CRISPR-Cas9 and raises concerns regarding pre-existing inflammatory microenvironment in patients, which may lead to adverse side effects during gene editing.

## RESULTS

### AAV delivery of CRISPR-Cas9 induces local inflammation

Before assessing the immune responses induced by the CRISPR-Cas9 system, we first examined the efficiency of AAV-delivered Cas9. AAV-SpCas9 was constructed as reported in Wu et al. with an sCMV promoter (*20*), but with a single sgRNA that has previously shown high efficiency in the retina (*3*) (Figure 1A). A dose of 5×10⁹ vg/eye is commonly used in experimental gene therapy studies (*21*). Therefore, we selected this dose for all our experiments, referred to as ‘standard dose’. However, one limitation of the dual AAV strategy is that achieving 5×10⁹ vg/eye for each component (Cas9 and sgRNA) requires a total dose of 1×10^10^ vg/eye. This higher viral load is necessary to maintain sufficient transgene expression from each AAV, but it also increases the risk of inflammation and immune responses. For this reason, we also evaluated this dose hereafter termed the ‘double dose’.

**Figure 1.**
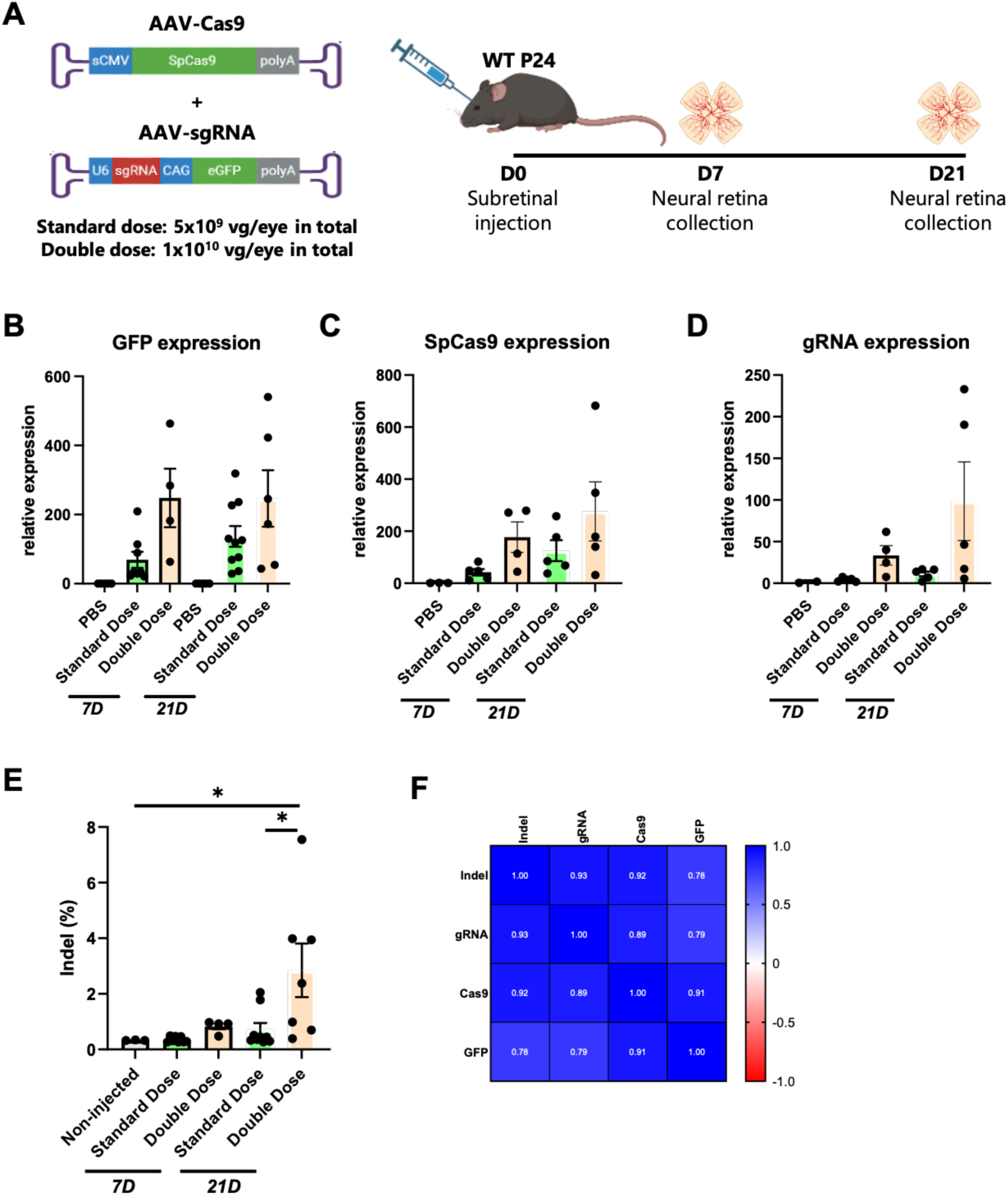
AAV-Cas9 is delivered subretinally and induces gene editing in mice. **(A)** Schematic of AAV cassette and delivery; **(B)-(D)** Expressions of transgenes including GFP **(B)**, Cas9 **(C)** and sgRNA **(D)**; **(E)** Indels in the whole neural retina after subretinal injection of AAV-Cas9 *in vivo.* Each dot represents neural retina isolated from a single mouse eye. Mean ± SEM. Ordinary one-way ANOVA test was used for multiple comparison; **(F)** Correlation matrix based on the transgene expressions and indels in mice receiving AAV injection 7 days and 21 days after. Pearson correlation (r) coefficients was used for correlation matrix.

Retinas from wild-type (WT) mice were collected after subretinal injection to investigate the impact of Cas9-AAV. The three transgenes (GFP, SpCas9 and sgRNA) were correctly expressed with increases over time (Figure 1B-D). We then quantified indels in the whole neural retina. Double dose AAV induced the highest indel levels 21 days after injection (Figure 1E). Transgene expression and editing efficiency correlated significantly (Figure 1F).

We then evaluated retinal thickness and the expression of inflammatory genes to assess the local inflammation. Retinal thickness measurement revealed no significant decrease across treated groups (Figure 2A). However, a decrease in thickness was observed in retinas that received double dose AAV. Thus, we analyzed the mRNA expression levels of inflammation-associated markers by droplet digital PCR (ddPCR). We selected CYBB, a marker for type 1 macrophages and phagocytosis (*22*); CD68, a marker for activated microglia/macrophages (*23*); CD18, important for leukocyte adhesion and transmigration (*24*); and CD11c, a marker of dendritic cells. All of these markers were significantly upregulated at 21 days post-injection in the double dose AAV-Cas9 group but not in the AAV-GFP control group. Although CD11b expression, a marker of macrophages and microglia (*25*), did not show significant changes, the same upward trend was observed in the double-dose group (Figure 2F). Principal component analysis (PCA) was done to visualize the difference among conditions based on the evaluated immune markers. There was no clear classification 7 days after the injection (Figure 2G) while 21 days after the injection, mice receiving double dose AAV were separated from other mice meaning the alternation of immune profiles (Figure 2H). Importantly, the immune responses induced by AAV-Cas9 at 21 days did not negatively correlate with indel levels (Figure S1). Taken together, these results show that double-dose AAV-Cas9 delivery triggers a local inflammatory response in WT retinas that is specific to CRISPR-Cas9, but does not compromise editing efficacy.

**Figure 2.**
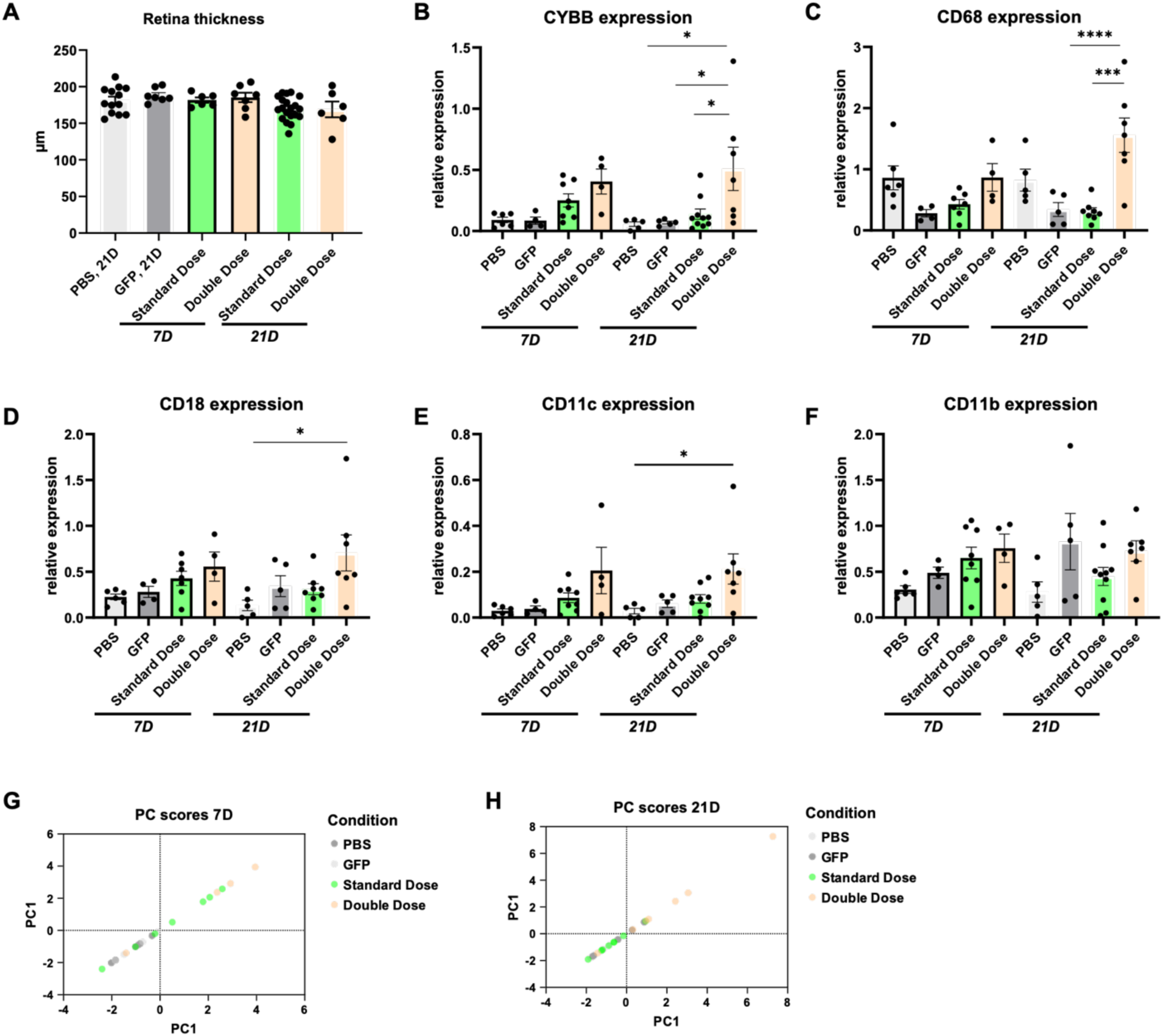
AAV-Cas9 induces local inflammation in mice. **(A)** Measurement of retinal thickness of mice receiving PBS, AAV-GFP (Standard dose: 5×10^9^ vg/eye) or AAV-Cas9 (Standard dose: 5×10^9^ vg/eye, or double dose: 1×10^10^ vg/eye); **(B)-(E)** Expressions of immune marker genes including CYBB **(B)**, CD68 **(C),** CD18 **(D),** CD11c **(E)** and CD11b **(F)**. Each dot represents neural retina isolated from a single mouse eye. Mean ± SEM. Ordinary one-way ANOVA test was used for multiple comparison. **(G)-(H)** Principle component analysis (PCA) based on the expressions of immune marker genes 7 days **(G)** or 21 days **(H)** after injection.

### AAV-Cas9 induced time- and dose-dependent immune responses

To provide a more comprehensive analysis of transcriptomic alterations in the retina after AAV-Cas9 delivery in WT mice, we performed bulk RNA sequencing of retinas injected with PBS, AAV-GFP standard dose, AAV-Cas9 standard or double dose. We then quantified differentially expressed genes (DEGs) of AAV-Cas9 compared to PBS or AAV-GFP at 7 and 21 days post-injection. We selected these two time points to capture the early innate immune activation (*26*) and the subsequent adaptive immune response (*13*).

Seven days after injection, AAV-GFP induced a minimal transcriptomic response with only 13 DEGs, whereas standard dose AAV-Cas9 led to 34 DEGs. At 21 days after injection, AAV-GFP continued to show minimal effects with 4 DEGs, while standard dose AAV-Cas9 induced 83 DEGs, suggesting a specific transcriptomic response against CRISPR/Cas9 (Figure 3A-B). Double dose AAV-Cas9 triggered a stronger response at both 7 and 21 days post-infection (Figure 3A-B), indicating a dose effect. The strongest transcriptomic responses were observed at 21 days post-injection, suggesting a prolonged impact of AAV-Cas9 on retinal gene expression.

**Figure 3.**
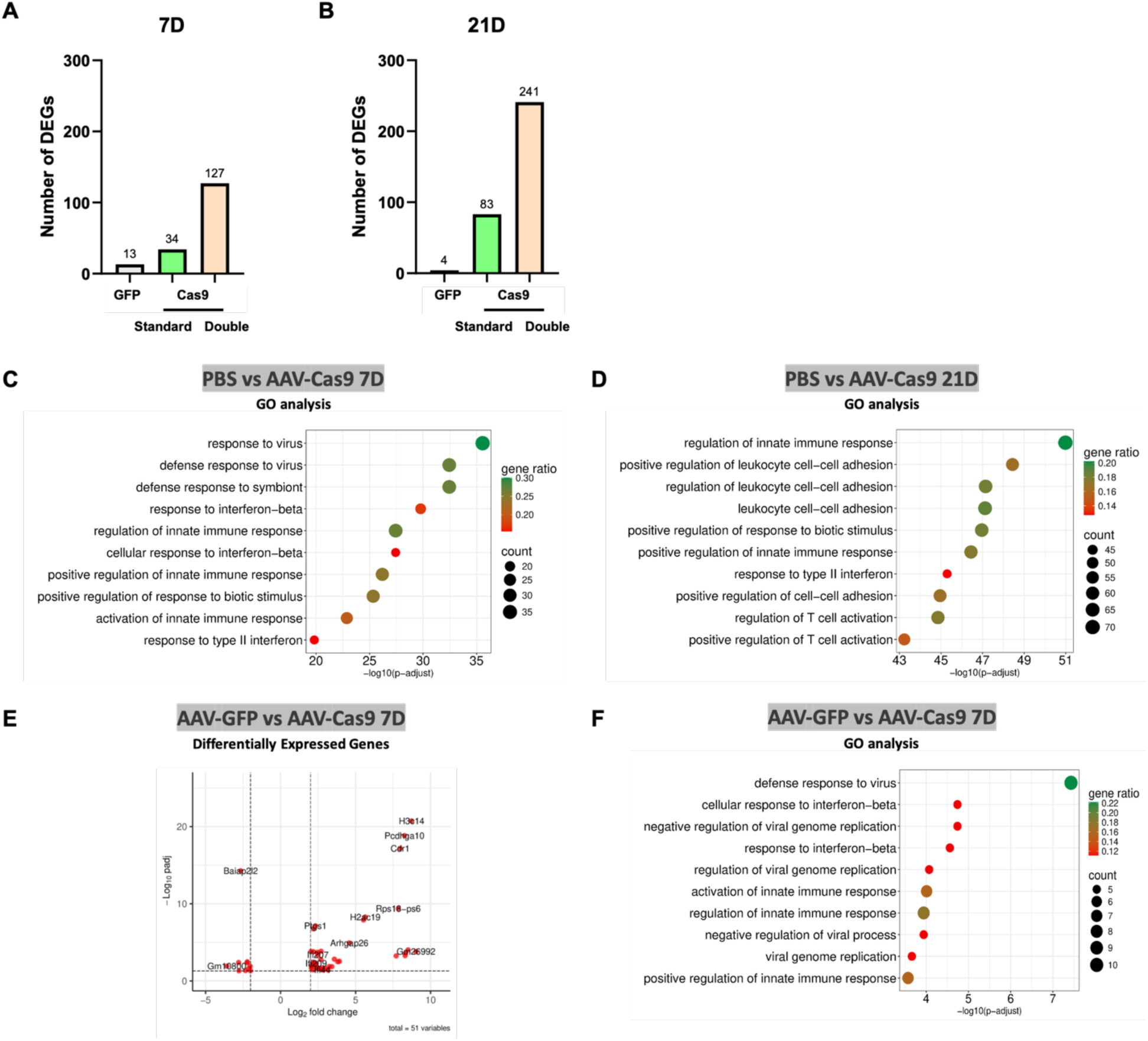
AAV-Cas9 induced time- and dose-dependent transcriptome alternation, specific to CRISPR-Cas9. (A)-(B) The numbers of differentially expressed genes (DEGs) in the retina receiving AAV-GFP (Standard dose: 5×10^9^ vg/eye) or AAV-Cas9 (Standard dose: 5×10^9^, or double dose: 1×10^10^ vg/eye) compared to PBS 7 days (A) and 21 days (B) after injection; (C-D) Pathway enrichment analysis in the retina receiving RNP compared to PBS 7 days (C) and 21 days (D). Log2FoldChange>2 or <-2. (E) Volcano plot showing DEGs in the retina receiving AAV-Cas9 (5×10^9^ vg/eye) compared to AAV-GFP (5×10^9^ vg/eye) 7 days after injection; (F) Pathway enrichment analysis in the retina receiving AAV-Cas9 (5×10^9^ vg/eye) compared to AAV-GFP (5×10^9^ vg/eye) 7 days after injection; Log2FoldChange>2 or <-2.

To determine which pathways were regulated following AAV-Cas9 injection, we performed Gene Ontology (GO) enrichment analysis comparing double dose AAV-Cas9 to PBS-injected eyes. At day 7, GO terms were dominated by antiviral and innate immune responses, with strong enrichment in pathways such as “response to virus” and “interferon response” (Figure 3C). By day 21, enrichment shifted toward pathways associated with immune-cell regulation, activation, and adaptive responses (Figure 3D). Notably, the presence of T-cell–related terms indicated a transition from an innate response to adaptive immune activation, demonstrating that AAV-Cas9 induced time- and dose-dependent immune responses.

To confirm the impact of CRISPR-Cas9 transgene on immune responses, we compared AAV-Cas9 standard dose with AAV-GFP. As shown in the volcano plot, we observed upregulation of interferon-stimulated genes in AAV-Cas9–treated retinas, including the interferon activated gene 211 (*Ifi211*) (Figure 3E). GO analysis of the DEGs between AAV-GFP and AAV-Cas9 confirmed enrichment in processes such as “defense response to virus” and “response to interferon-beta” (Figure 3F), suggesting that Cas9 provokes broader immune activation than GFP, which is known to be immunogenic (*27*).

### CRISPR/Cas9 RNP also induces local inflammation

Direct delivery of CRISPR-Cas9 protein complexed with its guide RNA, as naked RNP, can provide a safer option for *in vivo* gene editing than AAV delivery (*3*). However, the immune responses to RNP delivery have not yet been studied or compared to AAV-Cas9 delivery.

Similar to AAV-Cas9, WT mice received a subretinal injection of CRISPR-Cas9 RNP, and retinal samples were collected and analyzed at 7 and 21 days post-injection (Figure 4A). Genome editing showed significantly higher indel frequency at 7 days in the RNP-injected group compared to PBS controls, followed by a decline at 21 days (Figure 4B). This decline could be explained by the fact that RNP induced a significant decrease in retinal thickness at 21 days post-injection (Figure 4C).

**Figure 4.**
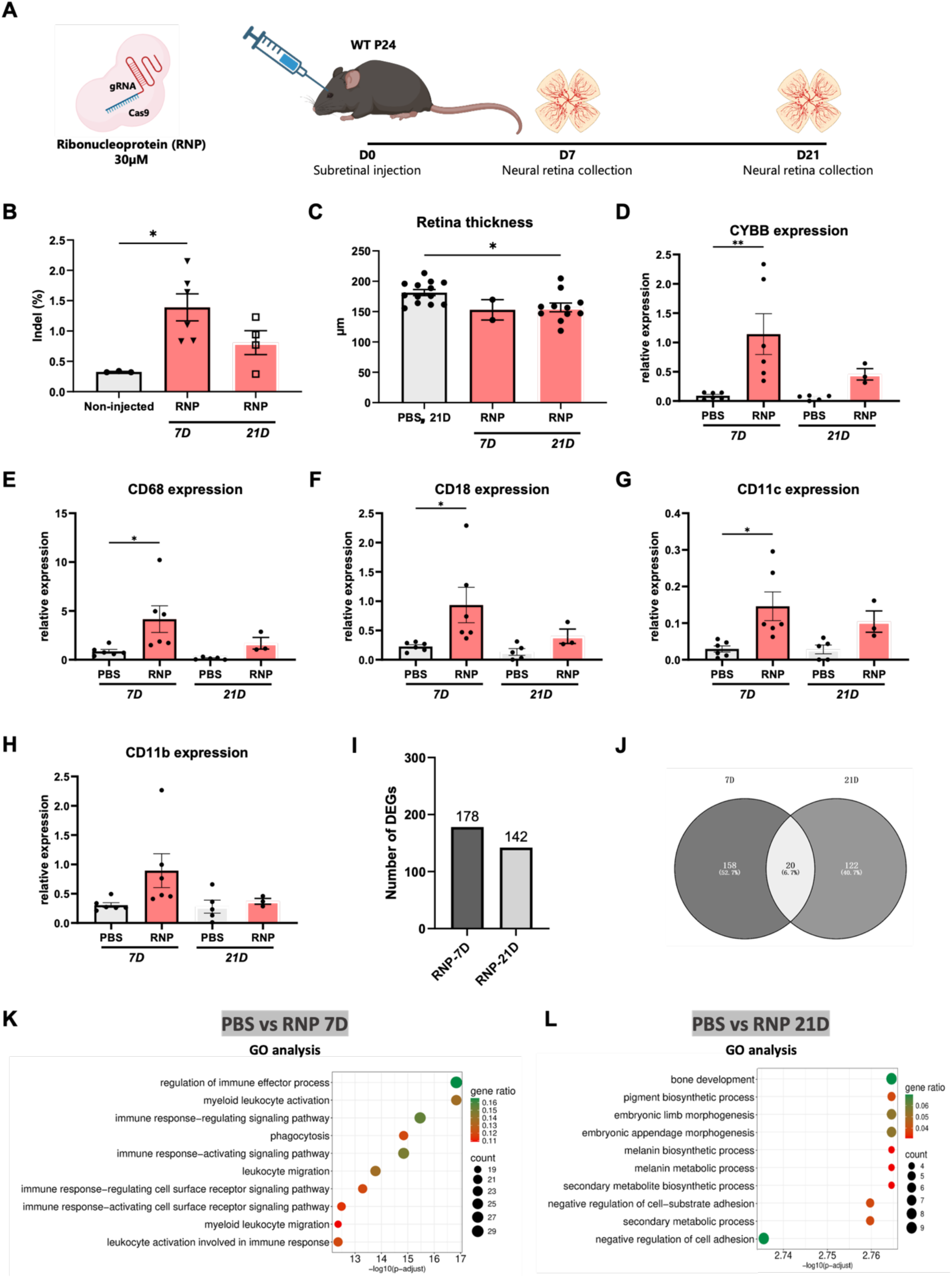
RNP induced gene editing and local inflammation in WT mice. **(A)** Schematic of RNP and delivery; **(B)** Indels in the neural retina after subretinal injection of RNP *in vivo;* **(C)** Measurement of retinal thickness of mice receiving PBS or RNP; **(D)-(G)** Expressions of immune marker genes including CYBB **(D)**, CD68 **(E)**, CD18 **(F),** CD11c **(G)** and CD11b **(H)**; **(I)** The numbers of DEGs in the retina receiving RNP compared to PBS 7 days and 21 days after injection. **(J)** Venn diagram showing the overlapped DEGs; **(K)-(L)** Pathway enrichment analysis in the retina receiving RNP compared to PBS 7 days **(K)** and 21 days **(L)** after injection. Each dot represents neural retina isolated from a single mouse eye. Mean ± SEM. Ordinary one-way ANOVA test was used for multiple comparison. Log2FoldChange>2 or <-2.

Analysis of inflammatory mRNA expression levels by ddPCR revealed a transient upregulation of CYBB, CD68, CD18 and CD11C at day 7, which declined by day 21 (Figure 4D to G). CD11b showed the same upward trend at day 7 (Figure 4H). A correlation analysis demonstrated correlations among immune markers, which do not correlate with editing efficacy (Figure S2). These results indicate that Cas9 RNP induced local inflammation 7 days after injection, which decreased 21 days after injection.

Transcriptomic analysis of neural retinas identified a higher number of DEGs at 7 days (n=178) compared to 21 days (n=142), with only 20 genes overlapping between the two time points (Figure 4I and J). GO enrichment analysis of DEGs confirmed that immune-related processes were highly enriched at 7 days, including regulation of immune effector processes and myeloid leukocyte activation (Figure 4K). By contrast, at 21 days, these immune-related pathways were no longer significantly enriched and were replaced by structural remodeling, such as pigment biosynthetic processes (Figure 4L). These results are consistent with transient activation of immune cells and innate immune signaling pathways, indicating that RNP-induced immune activation is short-lived.

### AAV-Cas9 and RNP delivery triggered distinct transcriptional and immune responses in the retina

To evaluate the immune response triggered by different genome editing delivery systems, we compared the expression of representative immune-related genes in the retina using bulk RNA sequencing. Expression of general immune cell markers such as CD45, H2-Eb1, and CD11b was upregulated in retinas treated with AAV-Cas9 at day 21 compared to PBS controls, indicating a persistent immune response. In contrast, RNP induced a transient upregulation of these markers at 7 days post-injection, demonstrating the short-lived immune response (Figure 5A). CD20, a marker for B cells, was not regulated (Figure 5B). Additionally, there was a trend toward increased expression of T cell markers in AAV-treated retinas at 21 days (Figure 5C), indicating the participation of adaptive cellular immune responses in AAV-Cas9 delivery.

**Figure 5.**
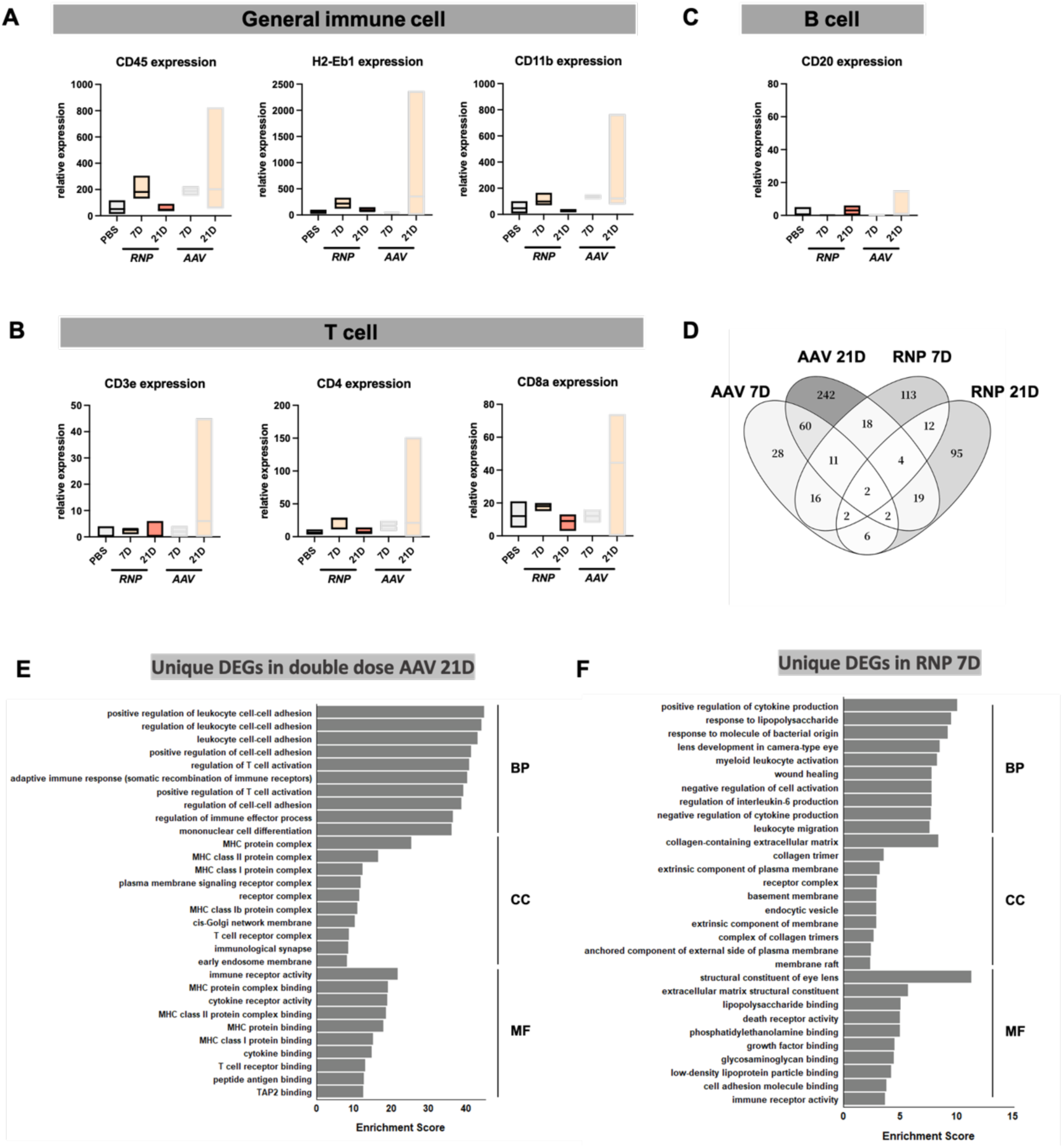
AAV-Cas9 and RNP delivery triggered distinct transcriptional and immune responses in the retina. **(A)-(C)** Expressions of immune marker genes including CYBB **(A)**, H2-Eb1 **(B)** and CD3e **(C)**; **(D)** Venn diagram showing the overlapped DEGs in the retina receiving RNP or double dose AAV-Cas9 compared to PBS 7 days and 21 days after injection. Each dot represents neural retina isolated from a single mouse eye. Mean ± SEM. Ordinary one-way ANOVA test was used for multiple comparison; **(E)-(F)** Pathway enrichment analysis based on unique DEGs identified in the retina receiving high dose AAV at 21 days **(E)** RNP at 7 days **(F)** after injection. Log2FoldChange>2 or <-2. BP: Biological Process; CC: Cellular Component; MF: Molecular Function. Enrichment Score represents (-log10(p-value)).

To further investigate the specificity of the responses elicited by each condition, we compared DEGs across AAV-Cas9 and Cas9 RNP. The Venn diagram showed that the majority of the regulated genes were not shared between the different conditions (Figure 5D). AAV-Cas9 at 21 days had the highest number of unique DEGs (n = 242), followed by RNP at 7 days (n = 113), which has been associated with higher inflammation in previous ddPCR analysis. The DEGs analysis demonstrated a clear difference between AAV-Cas9 and RNP delivery regarding activated immune pathways.

To gain further insight, we performed GO enrichment analysis of DEGs specific to double-dose AAV-Cas9 at 21 days or RNP at 7 days post-injection. Enrichment scores were calculated from three different GO domains: Biological Process (BP), Cellular Component (CC), and Molecular Function (MF). DEGs unique to AAV-Cas9 at 21 days showed enrichment in adaptive immune-related processes (Figure 5E), including regulation of T-cell activation, MHC complex and MHC protein receptor binding. These results suggest that prolonged expression of Cas9 via AAV delivery leads to antigen-specific adaptive immune response. In contrast, DEGs unique to RNP-treated retinas at day 7 were enriched in innate immune and structural remodeling pathways, such as “positive regulation of cytokine production”, “collagen-containing extracellular matrix” and “lipopolysaccharide binding” (Figure 5F). These results indicate that Cas9 RNP induced a transient innate inflammatory response but did not sustain adaptive immune activation.

### Pre-existing inflammation in the retinas of rd10 mice

As gene therapy is ultimately designed to treat patients with retinal degeneration, which may lead to pre-existing inflammation, pathological models can be valuable for assessing efficacy and safety profiles (*28*, *29*). We thus used a retinal degeneration murine model (rd10), with progressive photoreceptor degeneration and pre-existing retinal inflammation (*30*). The degeneration peaks in photoreceptor death at P24, which defined the time point for injecting all animals (*30*). As expected, retinal thickness was significantly reduced in rd10 mice compared to WT, reflecting photoreceptor degeneration (Figure S3A). Transcriptome analysis between WT and rd10 retinas in the absence of treatment also revealed substantial transcriptional reprogramming within rd10 retinas, including downregulation of photoreceptor-specific genes and upregulation of immune-related genes (Figure S3B). GO enrichment analysis confirmed these trends, showing enrichment in pathways involved in antigen processing and photoreceptor functions (Figure S3C).

### Enhanced inflammatory response in the degenerating rd10 retina

To assess the efficiency and safety of AAV-Cas9 and RNP-mediated gene editing in rd10 mice, we evaluated transgene expression, editing activity, and retinal integrity after subretinal injection. As in WT mouse, transgene expression levels of GFP, SpCas9 and sgRNA were higher at day 21 when injecting AAV-Cas9 (Figure S4A-C). At 7 and 21 days post-injection, indel rates were significantly higher in the double-dose AAV group compared to PBS control and other groups (Figure S4D), and this was correlated with transgene expressions (Figure S4E). Notably, the RNP group also showed measurable indel formation, though at a lower level than AAV (Figure S4D). While we did not detect a significant change in retinal thickness after CRISPR–Cas9 injection, a decreasing trend appeared in mice treated with double-dose AAV and RNP at day 21 (Figure S4F).

To assess the immune consequences of Cas9 delivery via AAV or RNP in the pathological model, we performed transcriptomic and GO enrichment analyses. Consistent with observations in WT mice, AAV-Cas9 induced the highest number of indels at 21 days, with an incredibly high number of DEGs (786 genes) (Figure 6A). In contrast, Cas9-RNP delivery resulted in more DEGs at 7 days (n = 196) than at 21 days (n = 60) (Figure 6A). Venn diagram analysis showed that DEGs were largely condition-specific, with minimal overlap across groups. The AAV-Cas9 21 days group displayed the greatest number of unique DEGs, followed by Cas9-RNP at 7 days (Figure S5A). GO enrichment analysis of DEGs exclusive to AAV-Cas9 at 21D revealed enrichment in immune-related biological processes, particularly those associated with T-cell activation and antigen processing (Figure S5B). In contrast, DEGs specific to Cas9-RNP at 7 days were enriched in processes linked to epithelial repair and extracellular matrix organization (Figure S5C). These results suggest that the trends of the immune responses induced by Cas9 delivery in rd10 mice were similar to those in WT mice.

**Figure 6.**
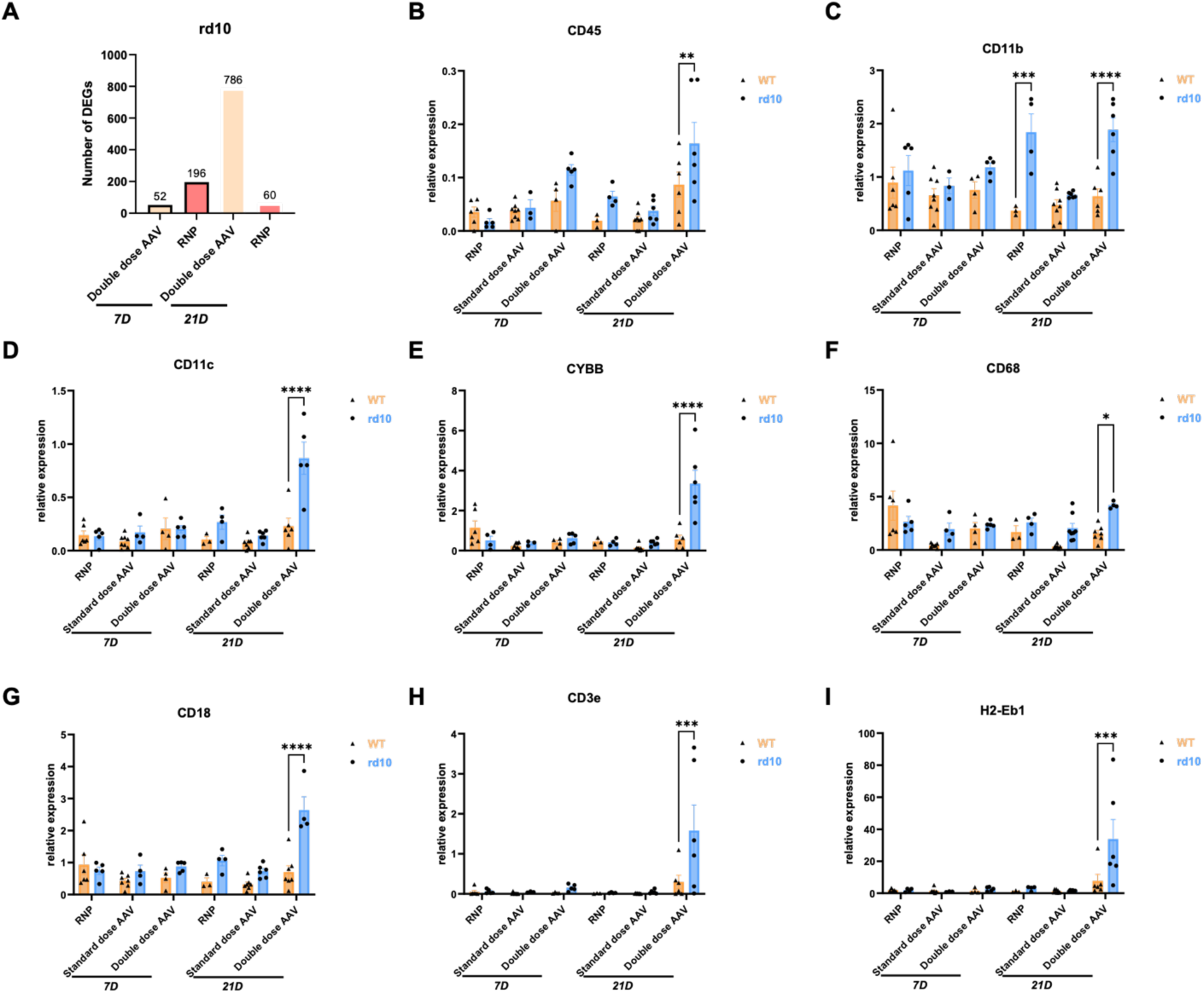
Enhanced inflammatory response was observed in rd10 mice compared to WT mice. **(A)** The numbers of DEGs in rd10 mice receiving double dose AAV or RNP compared to PBS 7 days and 21 days after injection; **(B)-(I)** Comparison of expression level of different immune markers including CD45 **(B)**, CD11b **(C)**, CD11c **(D)**, CYBB **(E)**, CD68 **(F)**, CD18 **(G)**, CD3e **(H)**, H2-Eb1 **(I)** from mice treated with RNP, standard dose or double dose AAV 7 days or 21 days after the injection. Each dot represents neural retina isolated from a single mouse eye. Mean ± SEM. Mann–Whitney test was used for pair comparison.

To determine whether retinal degeneration and basal inflammation contribute to the local immune response to CRISPR-Cas9, we compared inflammatory markers in WT and rd10 mice following subretinal delivery of AAV-Cas9 or Cas9-RNP. All markers studied were markedly higher in rd10 than in WT following double dose AAV delivery (Figure 6B-I). Notably, the adaptive immune marker CD3e, representing T-cell infiltration, was significantly upregulated in rd10 mice compared to WT at 21 days post-treatment with double-dose AAV (Figure 6H). This was consistent with a significant increase in the MHC II marker, H2-Eb1 (Figure 6I). To assess systemic immune responses, we measured antibodies against Cas9 and AAV in sera. However, no clear anti-Cas9 or anti-AAV antibodies were detected except in a few mice (Figure S6). For Cas9 RNP delivery, only CD11b was significantly higher in rd10 retinas compared to WT (Figure 6C). The exacerbated inflammatory signature in rd10 retinas suggests that retinal degeneration amplifies innate and adaptive immune responses to AAV-Cas9 delivery, with potential implications for gene therapy safety in disease contexts.

## DISCUSSION

CRISPR-Cas9 has emerged as a promising tool for precise gene editing with immense therapeutic value in the treatment of inherited disorders (*31*). However, due to its bacterial origin, the immunogenicity of CRISPR-Cas9 and the corresponding immune responses *in vivo* are still poorly understood. Adverse events reported in the CRISPR-Cas9-based clinical trial EDIT-101 in the eye have raised concerns about potential local immune consequences (*2*). To address this, we explored local inflammation induced by Cas9 delivered via AAV or RNP in murine retinas and found that both AAV-Cas9 and Cas9-RNP triggered distinct patterns of inflammation. Additionally, we observed exacerbated inflammation in a pathological model, underscoring the immune risks associated with CRISPR/Cas9-mediated gene editing.

The eye is considered an immune-privileged organ in which immune responses are typically restricted (*32*). Systemic delivery of Cas9 targeting organs such as the liver or muscle has been shown to induce T-cell responses capable of eliminating edited cells(*6*, *9*), and these responses are not reliably controlled by conventional immunosuppression (*2*, *9*). In the eye, studies have shown that innate and adaptive immune responses can be induced during the development of ocular gene therapy with AAV delivering various transgenes (*11*, *13*, *21*, *33*, *34*) but never specifically delivering Cas9. Thus, our study provides evidence that Cas9 can induce local inflammation in the retina, highlighting the potential risks of gene editing therapies even in immune-privileged sites. This risk is particularly relevant in the context of the dual AAV delivery strategy, which is often required due to the large size of the CRISPR-Cas9 system (*4*). Since this approach requires a higher vector dose to achieve sufficient amount editing, the dose-dependent immune response becomes more significant (*35*). Our results confirmed the exponential expressions of innate immune response markers like CD68 when doubling the dose. However, editing efficacy was relatively low across the whole neural retina, even at the highest tested dose. This highlights the need to improve CRISPR/Cas9 safety and efficacy in the retina, through constructs, promoter, and vector optimization, and route of delivery (*36*, *37*) as we confirmed that the dose is a limiting factor and cannot be infinitely increased. Strategies are emerging to incorporate CRISPR/Cas9 or even its derivative, base editors or prime editors, into a single AAV (*38–40*). However, the efficacy of those mini-constructs *in vivo* is yet currently limited (*41*).

Also, our results highlighted the major involvement of macrophages and microglia in the immune response against CRISPR-Cas9 regardless of the delivery method. Their activation following treatment point to a dynamic response consistent with roles in phagocytosis and antigen presentation, which was transient against Cas-RNP but persisted against AAV-Cas9. Those results are consistent with evidence showing that microglial activation can exert both neuroprotective and neurotoxic effects depending on the context and the duration of activation(*42*). Therefore, methods to regulate macrophage and microglial activity warrant investigation to mitigate the potential immune response in gene editing.

We also observed an increased CD3 marker and enrichment adaptive immune responses in AAV-treated groups indicating direct T-cell involvement. However, few mice developed detectable anti-Cas9 antibodies in the serum, suggesting minimal humoral responses, which is consistent with previous studies showing little antibody production in human and NHP subjects after ocular Cas9 delivery (*2*, *43*). A study comparing the serum and vitreous anti-Cas9 antibody levels in mice immunized with Cas9 systemically reports a lower presence of antibodies in vitreous fluid compared to the serum, suggesting restricted antibody access to the eye (*44*). Thus, investigation into the adaptive immune responses is needed to obtain the comprehensive understanding of activation and immigration of adaptive immune cells.

Beyond AAV, we also assessed local inflammation following naked RNP delivery of CRISPR/Cas9 which can provide transient presence of the gene editor, leading to fewer genotoxicity (*3*) and could minimize the potential inflammation (*45*). Nevertheless, we observed a significant reduction in retinal thickness, indicative of structural damage in this case, as well as a transient elevation in innate immune markers, demonstrating that RNP delivery still poses immune risks. In the future, non-viral vectors will be essential to enhance cellular uptake of RNPs and reduce the total administered dose, thereby improving the overall benefit/risk ratio (*46*).

We also observed different immune responses depending on the delivery method for CRISPR/Cas9, with distinct pathways involved. RNP induces a strong inflammatory response at 7 days, which is reduced by day 21, while AAV-Cas9 triggers both innate and adaptive immune responses that persist up to 21 days. Among these, unique immune responders can be identified and potentially targeted for the development of specialized immunosuppressive strategies, which could inform the optimal choice of vector. Additionally, biomarkers may be screened to represent the immune consequences of Cas9 delivery, facilitating the preclinical evaluation of immunogenicity in novel Cas9 variants (*47*).

Our findings also demonstrate that the underlying pathological condition of the retina significantly amplifies immune responses to gene editing interventions. In rd10 mice, upon AAV-Cas9 administration, innate and adaptive immune responses were markedly enhanced, surpassing those observed in WT mice. These results underscore the heightened immune vulnerability of degenerating tissues and highlight the importance of incorporating disease models in preclinical evaluations to more accurately assess safety and efficacy. Currently, murine models like Ai9-Rosa mice (*48*) or TIGER tdTomato mice (*49*) with reporters are preferred in exploring efficiency of gene editing tools but the corresponding immune consequences are often overlooked in these models that do not display pathology. This reinforces the value of developing pathological models that better reflect real-world conditions (*50*) which may facilitate more effective gene therapy strategies.

In conclusion, our study highlights the risk of CRISPR-Cas9-induced inflammation following ocular delivery via both viral and non-viral methods. The immune responses varied by vector type and time point, indicating complex immunogenic patterns: Cas9 RNP induces a short-term innate immune response while AAV-Cas9 induces innate and adaptive immune response. Transcriptomic analysis revealed differential pathway enrichment, opening avenues for further investigation. Finally, enhanced immune responses in pathological models underscore the need for safety evaluation of CRISPR-Cas9 strategies in vulnerable tissues.

## MATERIALS AND METHODS

### sgRNA design

sgRNA targeting mouse *Sag* gene (5’-AGACATGAAGAACTGCCAGG-3’) was used, as it had previously demonstrated efficiency in the neural retina (*3*). sgRNA was synthesized and purified using the GeneArt Precision gRNA Synthesis Kit (Invitrogen) as previously described (*3*). sgRNA eluted in water was aliquoted and stored at −80°C.

### Cas9 nuclease

For *in vivo* experiments, Streptococcus pyogenes Cas9 (SpCas9) nuclease with two nuclear localization sequences (NLS) (one on its N and one on its C terminal) was produced as previously described (*51*).

### RNP preparation and complexation

Ribonucleoproteins (RNPs) were prepared immediately before use as previously described (*3*). Briefly, SpCas9 proteins and sgRNAs were mixed at a molar ratio of 1:1 in a final buffer of 20 mM HEPES/200 mM KCL (pH7.4). The solution of SpCas9 and sgRNA was incubated at room temperature before use.

### AAV production

Plasmids were constructed according to Wu et al (*20*). The AAV.SpCas9 was constructed exactly as Wu et al. with an sCMV promoter. As for the AAV.sgRNA, sgA was replaced with our designed Sag sgRNA and sgB was removed and replaced with a CAG-EGFP reporter as previously described (*3*). The plasmids were purchased from VectorBuilder.

### Subretinal injections in mice

All animal experiments were realized in accordance with the NIH Guide for Care and Use of Laboratory Animals (National Academies Press, 2011). The protocols were approved by the Local Animal Ethics Committees and conducted in accordance with Directive 2010/63/EU of the European Parliament. The project was evaluated by the CEEA 05 (Ethical Committee in Animal Experimentation 05) and approved by the MESRI (“ministère de l’enseignement supérieur, de la recherche et de l’innovation”, France). The approval numbers of the projects from the animal facility are B-75-12-02 and C-75-12-02.

Two mouse models were used in this study: C57BL/6j, rd10/rd10 (rd10) mice carrying a mutation in *Pde6b* gene expressed by rods, leading to a dysfunctional phototransduction cascade and rod-cone dystrophy. Males and females were used indistinguishably.

Mice were anesthetized by isoflurane inhalation. Pupils were dilated and bilateral subretinal injections (dorso-temporal injections) of 1 μL were performed using a Hamilton syringe with a 33-gauge blunt needle (World Precision Instruments, Inc.) under an operating microscope (Leica Microsystems, Ltd.). Ophthalmic ointment (Fradexam) was applied after surgery. Eyes with extensive subretinal hemorrhage were excluded from the analysis. Animals were euthanized by CO2 inhalation and cervical dislocation.

### DNA/RNA extraction, quantification and reverse transcription

DNA and RNA were extracted from the whole neural retina of mouse eyes using Quick-DNA/RNA^TM^ Microprep Plus Kit (Zymo research) with the protocol provided. The concentration of DNA is measured with Nanodrop 2000. The quality and quantity of RNA were evaluated using BioAnalyzer 2100 (Agilent Technologies) with the RNA 6000 Nano Kit (Agilent Technologies). cDNA synthesis was performed following SuperScript® IV First-Strand cDNA Synthesis Reaction protocol.

### Quantification of indels

As described previously (*3*), PrimeSTAR GXL DNA polymerase (Takara) was used for PCR amplification on the targeted region, on the *Sag* gene (primer fw =5’-GCTCTGTGCGGTTACTGATC-3’; primer rv = 5’-AGTGGAAATAAGCTGGTAATTTGCA-3). The thermal cycler program for PCR was as follows: 98°C for 10 s, followed by 60°C for 15 s, and finally 68°C for 20 s, with 30 cycles in total. PCR samples were then purified using NucleoSpin PCR and gel kit (Macherey-Nagel) following the manufacturer’s instructions. Purified amplicons were verified by electrophoresis on a 2% agarose gel and then were sent for next generation sequencing (NGS) at the Massachusetts General Hospital DNA core facility. A minimum of 40,000 reads were generated per sample. Interleaved Fastq files were subsequently analyzed using CRISPResso2 (*52*). One point represents indels detected in the whole neural retina of one eye. Default analysis parameters were maintained. Editing percentages from the initial output figure representing modified reads were retained in accordance with the recommendations of CRISPResso2 developers.

### qPCR and ddPCR analysis

For qPCR, a mix of reaction solution is prepared with 10µL Master Mix Universel Power SYBR green (Applied Biosystems™): with 10ng cDNA and 10mM primers (Eurofin genomics) for one gene. Samples were run at duplicates with blank controls to confirm the absence of genomic DNA. QuantStudio 5 PCR System was used with the program as 10 min at 95°C, 30 s at 95°C and 1 min at 60°C for 40 cycles, 1 min at 60°C and 15 s at 95°C. 2^ΔΔCT was calculated based on the control condition as PBS injected group to represent the expression of target gene.

For ddPCR, 11 µL of ddPCR Supermix for probes (no dUTP) (Biorad), 1 µL from the primers/probes mix (the ratio of housekeeping gene and target gene was 1:1), 2µL of the cDNA sample from mice (4ng), and 6µL of water were added in one well. The droplets were generated by the Automated Droplet Generator (Bio-Rad). A PCR was proceeded by the C1000 Touch Thermal Cycler (Bio-Rad), with the program as 10 min at 95°C, 30 s at 94°C and 1 min at 58.1°C for 40 cycle, 10 min at 98°C, 12°C infinite hold. The results were obtained by QX 200 Droplet Reader (Bio-Rad) with the software QXManager for the analysis. Samples were run at duplicates with blank controls and copy numbers of the genes were obtained. Copy number of gene of interest was normalized with copy number of housekeeping gene and the ratio was multiplied by 100.

For both ddPCR and qPCR, one point represents the mRNA expression level of the tested gene in the whole neural retina of one eye. Primers used are listed in Table S1.

### RNA sequencing and analysis

RNA samples with RNA integrity number (RIN) over 7.0 were selected for RNA sequencing. 300 ng of RNA in 20 µL was prepared and bulk RNA sequencing was performed with NovaSeq X (Ilumina) and Novaseq X plus 10B reagent kit provided by the IGenSeq Genome Sequencing Platform in Paris Brain Institute. RNA sequencing analyses were performed as described in Couturier et al (*53*). Fastq files obtained from the sequencing were aligned using STAR (v2.7.9a) against the <organism> reference genome from Ensembl <version> (generated <date>), with option “--quantMode GeneCounts” to extract the raw counts for each gene, and all count files were concatenated into a single file. A sample file was created with the sample data including the conditions (<conditions>) and the replicate numbers.

The count file and the sample file were loaded in our in-house R Shiny application EYE DV seq. We added 1 to all counts in the count file to avoid any 0 read count errors. We then removed the genes having a total count < 10 across all samples. Finally, the DESeq2 (v1.40.2) analysis was performed comparing the groups defined in the ‘condition’ column, with ‘ <control>’ as control group and ‘<condition>’ as condition.

The results were then filtered for significant genes using a p-value < 0,05. An enrichment analysis was performed to study the implicated pathways, using the GO database (using the Bioconductor packages ‘<organism>’ (v 3.17.0) and ‘clusterProfiler’ (v 4.8.1))(*54*) and the figures were made with SRplot (*55*) and VENNY2.1(*56*).

### Enzyme-linked immunosorbent assay (ELISA)

96-well Maxisorp plates (ThermoFisher scientific) were coated with full AAV8 capsid (5×10^8^ vg/well) or SpCas9 (0.5 μg/well) diluted with coating buffer (0.84% NaHCO3, 0.356%Na2CO3, pH9.5) overnight at 4°C. The wells were emptied and washed with blocking buffer (PBS 1X-6% milk) before incubating the plate with blocking buffer at room temperature for 2 hours. Serial dilutions of primary antibodies (Anti-AAV8 antibody, Hum Immu; Anti-SpCas9 antibody, GenScript) or serum were prepared during the incubation. Primary antibodies were diluted with dilution buffer (PBS 1X-1% BSA) to ensure a gradient on the plate to calculate the standard curve for the experiment. Murine sera were diluted to 1/1000 with dilution buffer for AAV Cas9 ELISA. After the incubation, primary antibodies dilution and serum dilution were added in the wells and the plate was incubated for 1 hour in the incubator at 37°C after 3 times washing with washing buffer (PBS 1X-0.05% Tween 20). 1/4000 dilution of secondary antibodies (Goat Anti-Mouse IgG, Southern Biotech) was prepared with dilution buffer. Secondary antibodies were added into each well after washing 3 times and the plate was incubated for 1 hour in 37°C incubator. The TMB reagent and stop solution (Thermo Scientific) were placed at room temperature 30 min before the end of the previous incubation. When the last incubation ended, the plate was washed 3 times with washing buffer. TMB reagent was added and a blue color appeared. Stop solution was added to stop the reaction after 10 min. The plate was read at 450 nm to get the optical density of each sample.

### SD-OCT and fundus imaging

Spectral-domain optical coherence tomography (SD-OCT) was performed 7 days or 21 days post injections. For pupil dilation, 0.5% tropicamide (Mydriaticum, Thea) and 5% phenylephrine hydrochloride (Neosynephrine, Europhta) were added to both eyes. The animals were anesthetized by inhalation of isoflurane (Isorane, Axience) and placed in front of the SD-OCT imaging device (MICRON 5; Phoenix). The eyes were kept moist with eye gel (Lubrithal, Dechra) during the whole procedure. Images from the temporal side of the eye are shown. Image acquisitions were performed following the instruction provided by supplier. InSight software was used to process the images.

## Statistical analysis

Statistical analyses were performed with GraphPad Prism V10.0. Ordinary one-way ANOVA test was used for multiple comparison. Mann-Whitney test was used for pair comparison, Pearson correlation (r) coefficients was used for correlation matrix. p value <0.05: *, <0.01: **, <0.001: ***, <0.0001: ****.

## Data and code availability

The data that support the findings of the current study are available from the corresponding author upon reasonable request.

## Acknowledgments

The authors would like to thank Xavier Guillonneau and Frédéric Blond for providing suggestions on RNA-seq analysis and Lauriane Przegralek for the dosage of the RNA samples. The authors would also like to thank Melissa Desrosiers, Camille Robert and Augustin Darennes Degaugue for AAV production. J.P. was supported by grants from the Fondation pour la Recherche Médicale (FRM SPF201909009287) and the Foundation Fighting Blindness (PPA-0922-0840-INSERM). D.R. was supported by grants from China Scholarship Council (202108070132). This work was supported by grants from LABEX LIFESENSES (ANR-10-LABX-65), IHU FOReSIGHT (ANR-18-IAHU-01), AFM, Inserm, Sorbonne Université and Paris Ile-de-France Region (DIM C-Brains and BioConvS).

## Author contributions

J.P., D.R., S.F. and D.D. designed experiments. A.D.C. produced Cas9 proteins. J.P. and H.M. performed subretinal injections in mice. D.R. and L.V. dissected mice eyes and performed DNA/RNA extraction. L.V. prepared and analyzed NGS samples. D.R., Y.Q. and L.V. performed and analyzed ddPCR/qPCR. D.R. and T.V.M. performed and analyzed the RNA sequencing data. J.P. and D.R. optimized and analyzed OCT recordings. D.R., J.P., and D.D. designed the study and wrote the manuscript. D.A., J.P.C and S. F. provided scientific input and gave feedback on the manuscript.

