## Supplemental information for "AAV-delivered CRISPR-Cas9 elicits persistent retinal immune responses compared with transient responses to RNP"

Supplemental results:

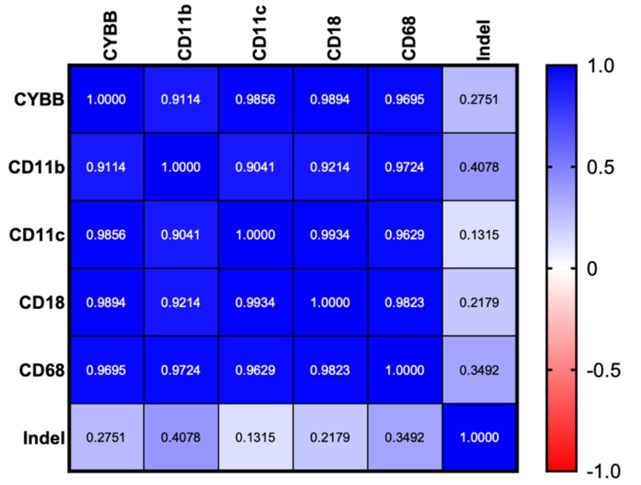


**Figure S1. Correlation analysis between local immune marker expressions and indels in WT mice receiving double dose AAV 21 days post-injection.** Pearson correlation (r) coefficients was used for correlation matrix.

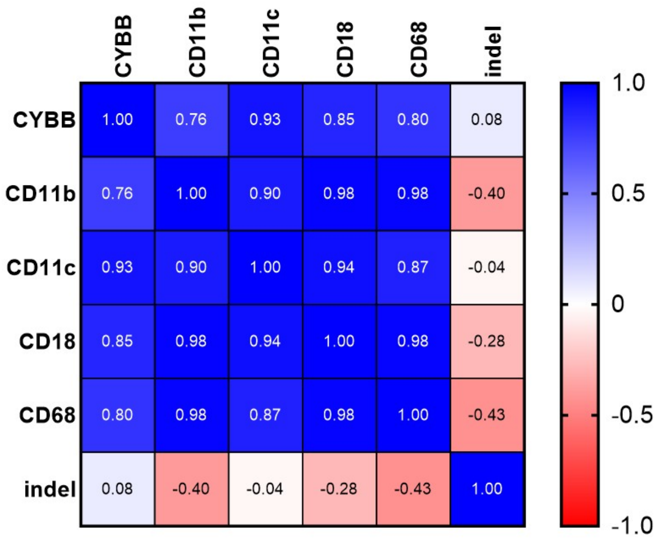


**Figure S2. Correlation analysis between local immune marker expressions and indels in WT mice receiving RNP 7 day after injection.** Pearson correlation (r) coefficients was used for correlation matrix.

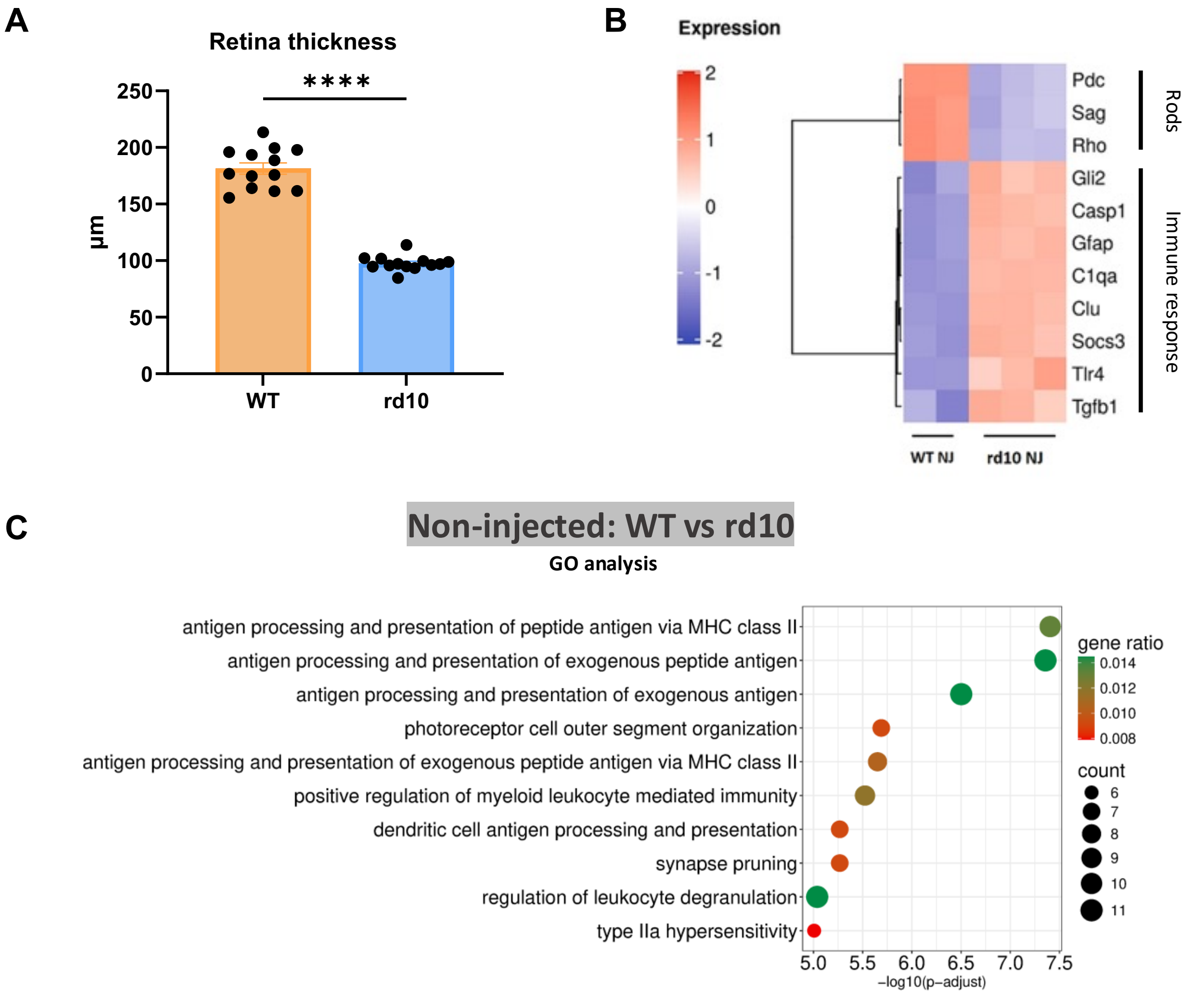


**Figure S3. Pre-existing inflammation was observed in the pathological model compared to WT mice with non-injected treatment (NJ).** (A) Expression matrix showing genes related to rods and immune response; (B) Measurement of retinal thickness of NJ WT and rd10 mice ; (C) Volcano plot and pathway enrichment plot between WT and rd10 mice. Each dot represents neural retina isolated from a single mouse eye at 45 days after birth. Mean ± SEM. Mann–Whitney test was used for pair comparison.

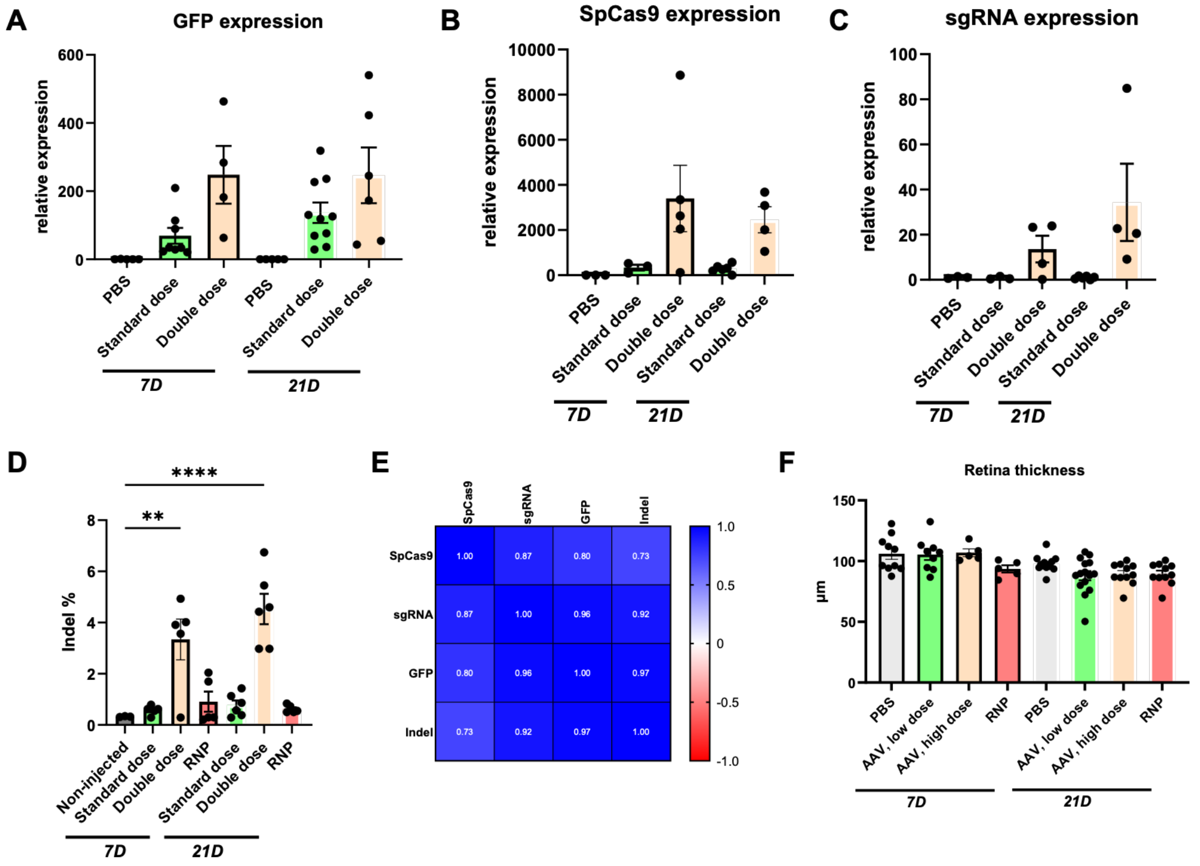


**Figure S4. AAV or RNP was delivered into rd10 mice for gene editing and induced inflammation.** (A)-(C) Expressions of transgenes including GFP (A), Cas9 (B) and sgRNA (C); (D) Indels in the neural retina after subretinal injection of AAV-Cas9 in vivo; (E) Correlation matrix based on the transgene expressions and indels in mice receiving AAV injection 7 days; (F) Measurement of retinal thickness of each group. Each dot represents neural retina isolated from a single mouse eye. Mean ± SEM. Ordinary one-way ANOVA test was used for multiple comparison.

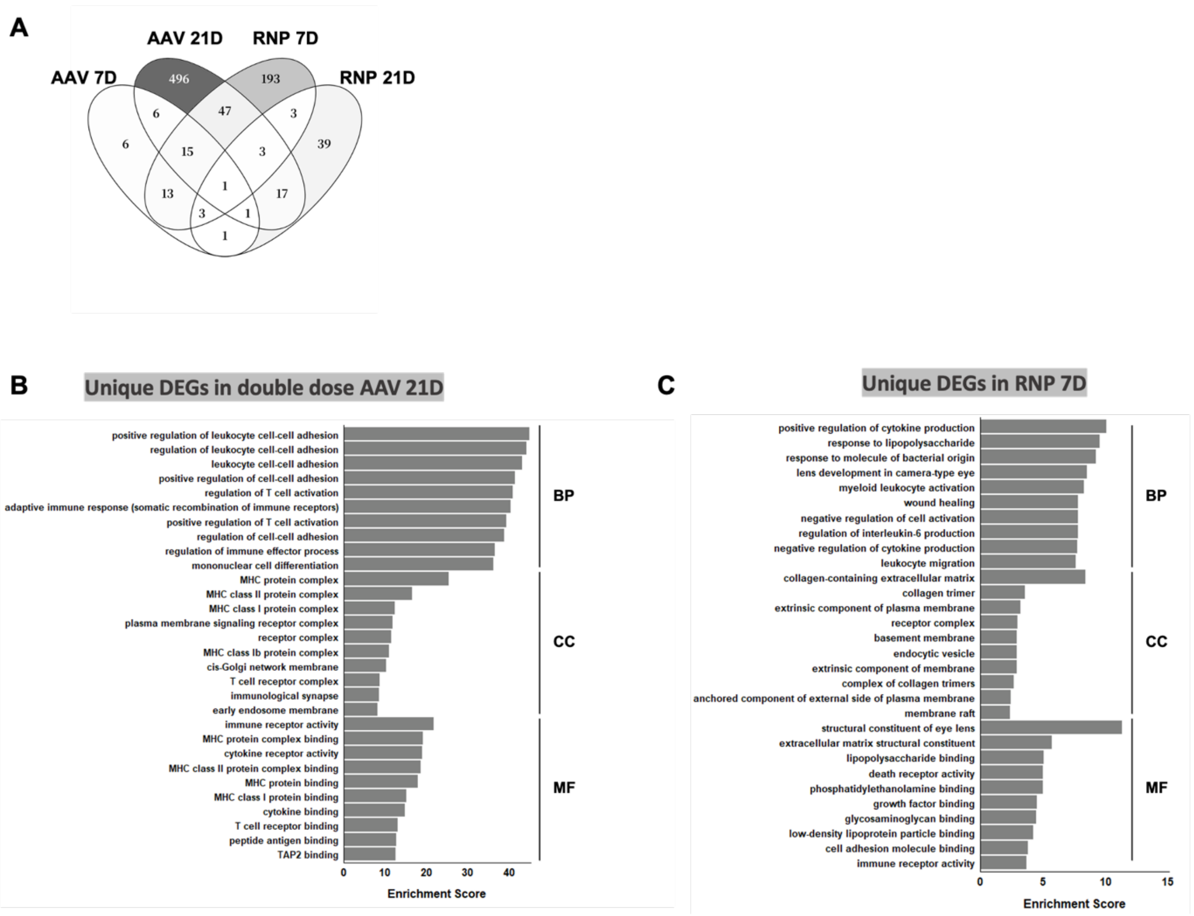


**Figure S5. AAV-Cas9 and RNP delivery triggered distinct transcriptional and immune responses in the retina.** (A) Venn diagram showing the overlapped DEGs; Pathway enrichment analysis in rd10 mouse based on unique DEGs identified in the retina receiving double dose AAV at 21 days (B) RNP at 7 days (C) after injection. Log2FoldChange>2 or <-2. BP: Biological Process; CC: Cellular Component; MF: Molecular Function. Enrichment Score represents (-log10(p-value)).

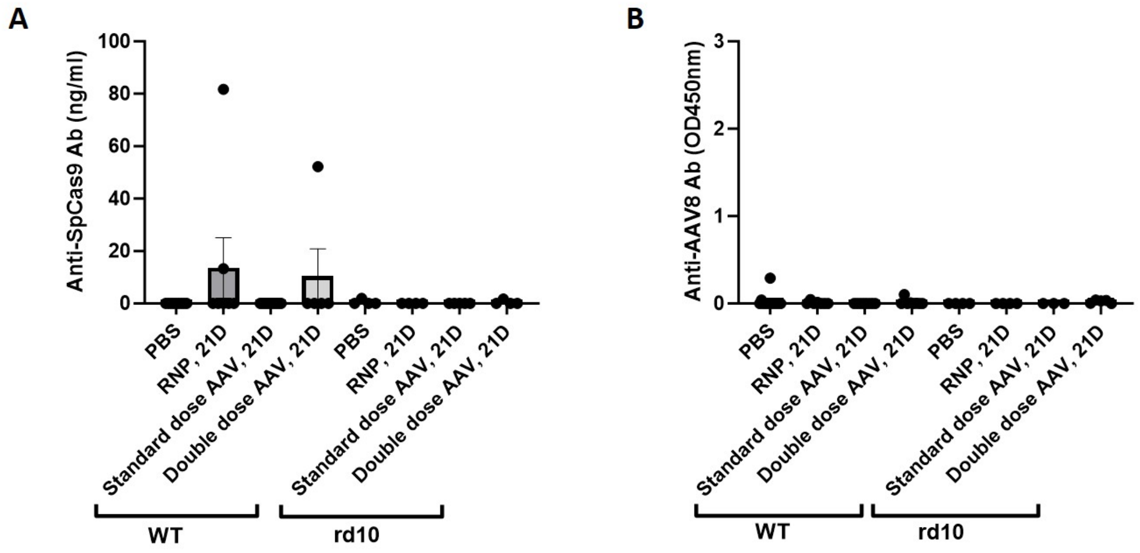


**Figure S6. Few mice produced anti-Cas9 antibodies while no mice produced anti-AAV antibodies.** (A)-(B) Evaluation of anti-Cas9 antibody (A) and anti-AAV antibody (B) in mice receiving PBS, AAV and RNP in WT and rd10 mice. Each dot represents serum from one mouse. Mean ± SEM. Mann–Whitney test was used for pair comparison.

### Table S1. Primers used for the RT-qPCR and dd-PCR

| Gene Name | Primer Name | Sequence (5′ → 3′) | Probe Name | Sequence (5′ → 3′) | Purpose |
|---|---|---|---|---|---|
| P0 | P0-F | CTCCAAGCAGATGCAGCAGA | P0-Probe | CCGTGGTGCTGATGGGCAAGAA | ddPCR |
| | P0-R | ATAGCCTTGCGCATCATGGT | | | |
| GFP | GFP-F | ACGTCTATATCATGGCCGAC | GFP-Probe | ACGGCCCCGTGCTGCTGCCC | ddPCR |
| | GFP-R | GTGCTCAGGTAGTGGTTGTC | | | |
| CYBB | CYBB-F | TCGAAAACTCCTTGGGTCAG | CYBB-Probe | CGGGCACCTGCAGCCTGCCTGAATT | ddPCR |
| | CYBB-R | TCTTCGAATCCTTGTCGAGC | | | |
| CD68 | CD68-F | CTAGGACCGCTTATAGCCCA | CD68-Probe | CCACAGGCCCACCACCACCAGTCAC | ddPCR |
| | CD68-R | ATGAGTGACAGTTGTGGGTC | | | |
| CD18 | CD18-F | GACTGAGAGGGGGACATGCT | CD18-Probe | TGCTGTGTCCCAGGAATGCACCAAGTACAA | ddPCR |
| | CD18-R | CTGGATACAGTCCCGGCAA | | | |
| CD11c | CD11c-F | GAGCTGTACCTGGATAGCCT | CD11c-Probe | CGGTGCTGAGTTCGGACACAGTGTGC | ddPCR |
| | CD11c-R | CAACCACCACCCAGGAACT | | | |
| CD11b | CD11b-F | GCTACGTAATTGGGGTGGG | CD11b-Probe | TGCCTTCAACAAACCACAGTCCCGCA | ddPCR |
| | CD11b-R | TTAGATGCGATGGTGTCGAG | | | |
| CD45 | CD45-F | GAGAGTAGGAAACTTGCTCCC | CD45-Probe | TGGCCTTTGGATTTGCCCTTCTGGACACA | ddPCR |
| | CD45-R | CCAAACATGGCAGCATCAC | | | |
| CD3e | CD3e-F | GCCATCTTGGTAGAGAGAGC | CD3e-Probe | TCTGGGGCATCCTGTGCCTCAGCCT | ddPCR |
| | CD3e-R | TCAATGTTCTCGGCATCGTC | | | |
| H2-Eb1 | H2-Eb1-F | CTGGTCCGAAATGGAGACTG | H2-Eb1-Probe | ACCTGCCAGGTGGAGCATCCCAGCC | ddPCR |
| | H2-Eb1-R | TGTGCTTTCCACTCGACC | | | |
| GAPDH | GAPDH-F | TTCACCACCATGGAGAAGGC | / | | qPCR |
| | GAPDH-R | CCC | | | |
| SpCas9 | SpCas9-F | AACAGCCGCGAGAGAATGAA | / | | qPCR |
| | SpCas9-R | CACGGGGTGTTCTTTCAGGA | | | |
| gRNA | gRNA-F | GACAAGCCCCTGAACCTCTC | / | | qPCR |
| | gRNA-R | TGTTGTTGGTCACGGTCACA | | | |
